# Exploiting outer-membrane protein promiscuity to induce transient collateral sensitivity via efflux pump competition

**DOI:** 10.64898/2026.05.12.724739

**Authors:** Pablo Catalán, Saúl Ares, Beatriz Pascual-Escudero, Ioannis Kontos, Pablo Laborda, José Ramón Valverde, José L. Martínez, Sara Hernando-Amado

## Abstract

In the fight against antimicrobial resistance, the identification of robust collateral sensitivity (CS) patterns could form the basis of successful sequential and combinatorial antimicrobial therapies. While most of the focus has been devoted to study stable CS due to mutations, CS can be transiently induced in *Pseudomonas aeruginosa* using dequalinium chloride (DC), which increases the expression of the efflux operon *mexCD-oprJ*. This leads to transient CS to the aminoglycoside tobramycin, whose molecular mechanism remains unclear. Using a combination of experimental results, mechanistic mathematical models and structural simulations, we show that the inactivation of NfxB by DC not only increases *mexCD* expression but also reduces the effective amount of the MexXY-OprM aminoglycosides efflux pump. Our data suggest that, under conditions of high MexCD production, this efflux pump can outcompete MexXY for the outer-membrane protein OprM, thereby inducing CS to tobramycin. Bayesian fits accurately reproduce measured *mexCD* expression and minimal inhibitory concentration (MIC) shifts across DC doses. Model predictions were validated in *P. aeruginosa* PA14 mutants lacking functional outer-membrane channels. Loss of OprJ preserves DC-induced CS to tobramycin, whereas loss of OprM abolishes it. These results identify competition for OprM as the proximal cause of DC-induced, transient CS and provide a generalizable framework for eliciting CS by perturbing shared components of multidrug efflux systems. This strategy suggests immediate avenues to enhance aminoglycoside efficacy while minimizing selection for stable resistance.

## 1 Introduction

Bacterial antimicrobial resistance (AMR) rates continue to increase worldwide ^1^, posing a major public health threat^2^. The opportunistic ESKAPE pathogen *Pseudomonas aeruginosa* alone was associated with more than 550,000 deaths in 2019 (44,000 in Europe)^3^, largely through nosocomial infections and chronic infections in people with cystic fibrosis (pwCF) or chronic obstructive pulmonary disease ^4,5.^ This pathogen exhibits low intrinsic susceptibility to many antibiotics ^6,7^ and readily acquires additional resistance through mutations ^8,9.^ Consequently, substantial efforts are underway to identify new strategies to combat AMR in this pathogen ^10^.

One phenomenon that has attracted increasing attention is collateral sensitivity (CS), whereby resistance to one antibiotic results in increased susceptibility to another ^10^. Although a few cases of CS associated with the acquisition of AMR genes by horizontal gene transfer have been reported ^11,12,^ most studies have focused on stable mutation-driven CS ^10,13-21.^ Despite the therapeutic potential of CS-based strategies, they still need to be clinically implemented. One major challenge is that distinct CS patterns can arise not only across different genetic backgrounds (i.e. pre-existing resistant mutants) but even among replicate populations of a single strain evolved under the same drug ^22-25^. Crucially, multiple *P. aeruginosa* isolates can coexist in pwCF undergoing antimicrobial treatment ^8,9^,26, and they may evolve diverging CS patterns when acquiring resistance to the same drug. Although such clonal variability can limit the clinical exploitation of CS, we have previously reported robust CS patterns with clinical potential against *P. aeruginosa*. In particular, the evolution of resistance to ciprofloxacin leads to phenotypic convergence toward CS to the aminoglycoside tobramycin. This phenotype is especially pronounced when associated with loss-of-function mutations in *nfxB*, which encodes a repressor of the RND ciprofloxacin efflux pump MexCD-OprJ ^17,27,28^.

RND efflux pumps are tripartite complexes composed of an inner-membrane transporter (the efflux pump itself), a periplasmic adaptor protein, and an outer-membrane channel. The genes encoding the inner-membrane and periplasmic components always form an operon; however, in some cases (e.g., *mexXY*) ^29^, this operon does not include the gene encoding the outer-membrane protein. Consequently, outer-membrane proteins can constrain the effective number and type of efflux pumps and exhibit some degree of promiscuity. For instance, the *P. aeruginosa* tobramycin efflux pump MexXY binds to the outer-membrane protein OprM, encoded in the *mexAB-oprM* operon ^30^, and the ciprofloxacin efflux pump MexCD, which associates with the outer-membrane protein OprJ, can also use OprM when overproduced ^31^.

In previous works we showed that evolutionary trajectories under ciprofloxacin selection are contingent on the genomic background and that ciprofloxacin exposure does not always result in overproduction of MexCD-OprJ through loss-of-function mutations in *nfxB* ^19,20.^ We therefore sought to induce CS to tobramycin while bypassing the necessity to acquire ciprofloxacin-resistance mutations. To this end, we exposed different isogenic mutants and clinical isolates of *P. aeruginosa* to dequalinium chloride (DC), an antiseptic that induces *mexCD-oprJ* expression. Transient overproduction of this efflux pump causes transient resistance to ciprofloxacin and increases susceptibility to the aminoglycoside tobramycin, mirroring the phenotype associated with loss-of-function mutations in *nfxB* ^27,32^. Moreover, treatment with the combinations DC-tobramycin resulted in extinction of the *P. aeruginosa* strains analyzed, suggesting that induction of CS may represent a promising strategy for designing effective antibiotic therapies against this priority pathogen. Despite this, the underlying molecular mechanism of transient CS remained unclear. Oure hypothesis was that the promiscuity of the outer-membrane protein OprM previously proposed may generate competition between the MexCD and MexXY efflux pumps. Such competition would increase the abundance of MexCD-containing efflux pumps while reducing the formation of MexXY-OprM complexes, ultimately increasing susceptibility to tobramycin.

In this study, we test our hypothesis by combining mathematical modelling, Bayesian inference, structural simulations and experiments using mutants lacking functional outer-membrane channels. We show that: *i*) DC induces *mexCD* expression by binding to its repressor NfxB; *ii*) MexCD competes with MexXY for OprM, thereby reducing the formation of functional tobramycin-exporting MexXY-OprM efflux pumps.

Taken together, our results provide mechanistic insight into how competition between efflux pumps can cause transient CS in *P. aeruginosa*, a phenomenon that could potentially be exploited to improve therapeutic strategies against this pathogen. More broadly, our findings offer a general framework for understanding promiscuity-driven CS patterns in other pathogens possessing orthologs of these efflux systems.

## 2 Results

### 2.1 *mexCD* is overexpressed by the use of dequalinium chloride

We first examined whether DC induces expression of the *mexCD* operon. Under basal conditions, *mexCD* is constitutively repressed by NfxB, a TetR-family transcriptional regulator ^33^. Although no crystal structure is currently available for NfxB, *in silico* structural models (see Materials and Methods) indicate strong similarity to the transcriptional regulator RamR. One key difference is that NfxB functions as a tetramer ^34^, whereas RamR is a dimer. RamR is known to bind DC, and this interaction reduces its affinity for DNA ^35^. Thus, we hypothesized that DC might inhibit NfxB in an analogous manner by directly binding to it.

To explore this possibility, we first estimated the binding affinity of DC for RamR using three independent binding-affinity prediction tools and published crystal structures ^35^. Docking simulations of DC using a SwissModel homology model of NfxB ^36^ and AutoDock Vina generated several energetically favorable binding poses (Figure 1A). Evaluation of their predicted affinities (Supplementary Table 1; see Materials and Methods) yielded consistent results: in nine out of ten poses, DC was predicted to bind NfxB with higher affinity than RamR. These results are consistent with the hypothesis that DC can bind NfxB and potentially interfere with its repression of *mexCD*. Docking analyses consistently predicted preferential binding of DC within the NfxB DNA-binding pocket, oriented parallel to Trp 115. The alignment of aromatic rings suggests π-π stacking interactions, similar to those obtained between DC and RamR residue Phe 155. In both cases, DC binds deeply within the DNA-binding channel, suggesting that this interaction could interfere with NfxB-DNA interactions.

**Figure 1.**
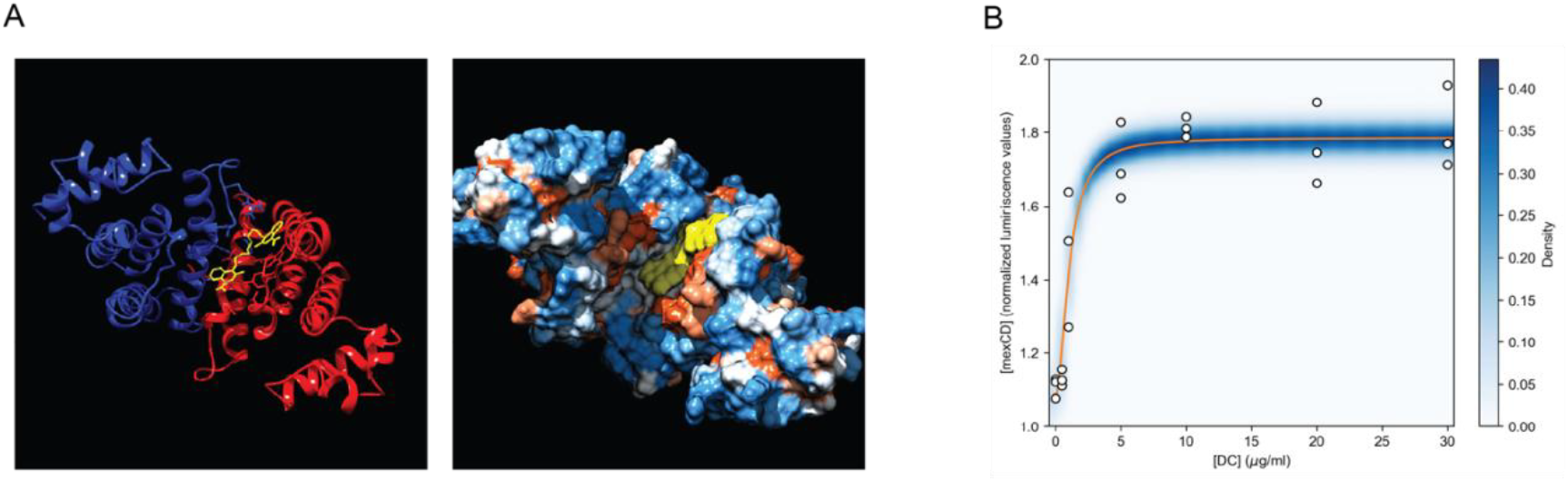
Structural and functional analysis of NfxB inhibition by dequalinium chloride. **A**. Ribbons and space-fill models of NfxB bound to dequalinium chloride (DC). Left: DC is shown in yellow, and NfxB subunits are shown as ribbons in blue and red. For clarity, only a dimer is shown, with the DNA binding groove facing the viewer. Right: surface model of two NfxB subunits (colored by hydrophobicity: blue hydrophilic, orange hydrophobic) bound to DC. **B**. *mexCD* expression as a function of DC. Orange line is equation (1), with *a*=1.79, *b*=0.8, *c*=1.67, obtained from linear regression, *R*^*2*^=0.90, which also coincides with the average values for the parameters obtained through Bayesian estimation. The blue shades represent the density of the posterior predictive distribution (i.e. the probability that *mexCD* expression has a certain value given a value of DC concentration, according to the posterior probability distribution of the parameters *a, b, c*, see Methods). Data (open circles) from ^32^.

Assuming that one molecule of DC binds to each NfxB dimer, and given that NfxB functions as a tetramer, we hypothesized that two DC molecules are required to fully inhibit NfxB activity. Based on this assumption, we derived a mechanistic mathematical model of the binding between DC and NfxB, and studied its effect on the expression of *mexCD* (see Supplementary Note 1), obtaining the following formula:

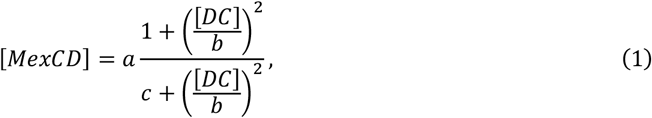

where [*MexCD*] and [*DC*] are the concentrations of MexCD and DC, respectively, and *a, b*, and *c* are unknown positive parameters. Using Bayesian estimation (see Materials and Methods), we obtained a posterior probability distribution for these parameters, combining equation (1) with previous *mexCD* expression data. The resulting fit accurately reproduces the observed expression dynamics (Figure 1B) and, together with our affinity predictions, supports a model in which DC binding inhibits NfxB and thereby induces *mexCD* expression.

### 2.2 OprM forms stable complexes with MexCD

OprM is an outer-membrane channel capable of interacting with multiple efflux systems. Although encoded in the *mexAB–oprM* operon and forming the MexAB–OprM efflux pump, it also partners with MexXY, whose operon lacks an outer-membrane protein ^37^. Moreover, mutants lacking OprJ (and thus, the canonical MexCD–OprJ efflux pump) remain resistant to ciprofloxacin, consistent with MexCD recruiting the alternative outer-membrane channel OprM to assemble a functional efflux complex ^31^.

To test the structural feasibility of this promiscuity, we performed molecular dynamics simulations of the relevant complexes. As a reference, we first simulated the cognate MexAB-OprM complex (PDB:6IOK). The complex remained assembled throughout the trajectory, and its overall structure was preserved, validating our simulation approach. We next assembled models of MexC (Fig. 2A) and MexD (Fig. 2B) into a MexCD complex (Fig. 2C), which was subsequently combined with OprJ (PDB:5AZS) to generate a MexCD–OprJ complex (Fig. 2D). During simulation, the complex remained structurally stable. Minor conformation changes were observed, including slight opening of some MexC arms distal to the MexCD interface and a modest bend at the MexC–OprJ junction. However, both interfaces were preserved throughout the trajectory. These deviations may reflect the absence of cellular envelope constraints in the simulation environment. We then assessed whether MexCD could form a stable complex with OprM. MexCD–OprM models (Fig. 2E) were constructed using OprM from PDB:6IOK, following analogous assembly procedures. Fully atomistic simulations at this scale required simulation boxes containing approximately 15 million atoms, resulting in substantial computational demands. To evaluate structural stability across extended timescales, we therefore performed both full-atom (5 ns) and coarse-grained simulations (100 ns). In the latter approach, the system size was reduced to approximately 0.75 million particles, accelerating computation and enabling longer simulation times while yielding results consistent with equivalent atomistic models. In these simulations, the MexCD-OprM complex remained assembled and maintained stable interfacial interactions. MexD appeared to press slightly into MexC, a movement that may be constrained *in vivo* by envelope architecture. This configuration supports a role for the MexC periplasmic arms in stabilizing MexD within the tripartite assembly. Overall, our results suggest structural compatibility between MexCD and OprM and provide mechanistic support for the feasibility of MexCD-OprM complex formation, which would cause a decreased amount of MexXY-OprM and, therefore, transient CS to tobramycin.

**Figure 2.**
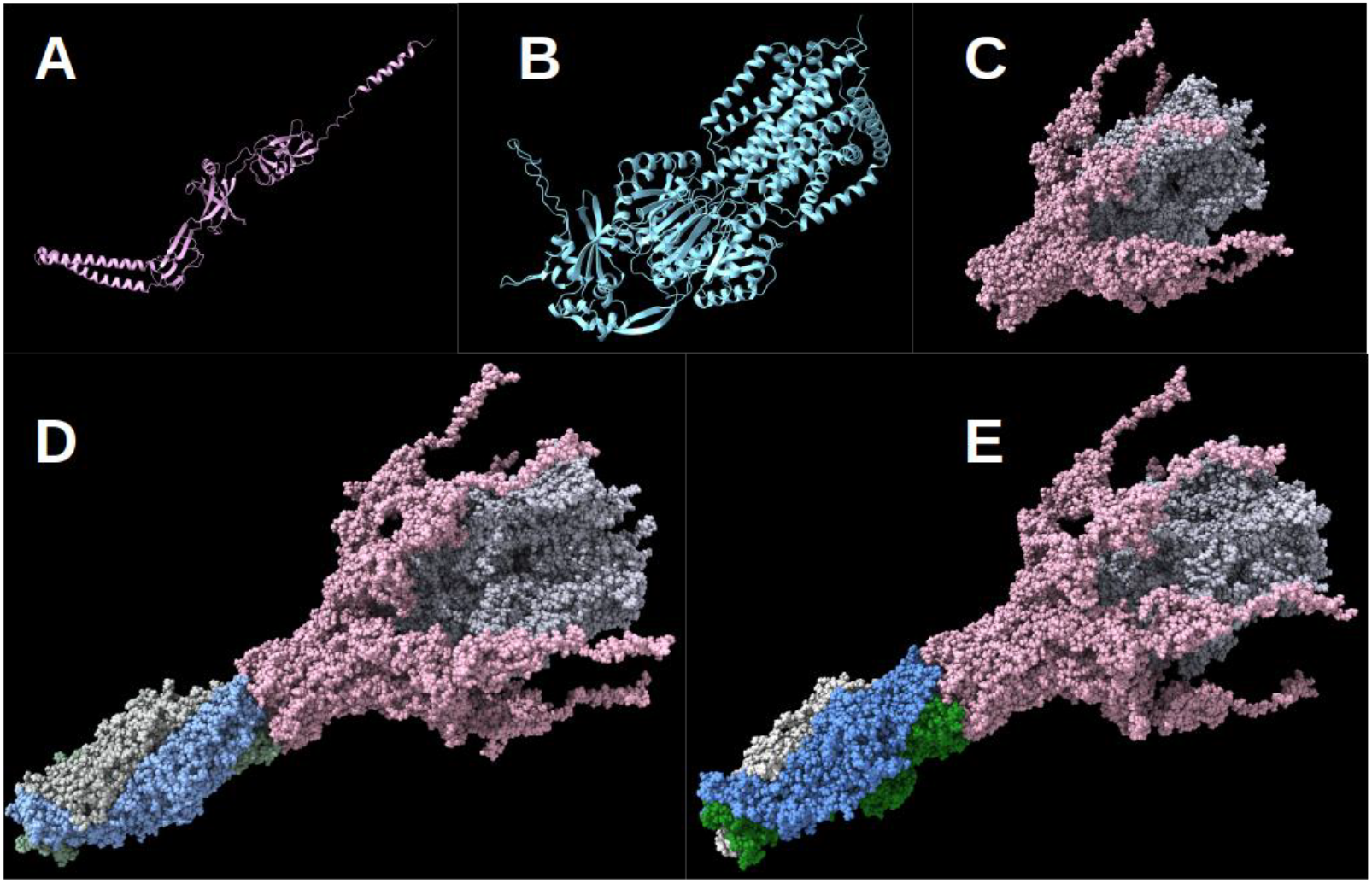
Incremental modeling of the MexCD complex with OprJ and OprM. A: Predicted model of MexC. B: Predicted model of MexD. C: Predicted model of the MexCD complex. D: Predicted model of the MexCD-OprJ complex. E: Predicted model of the MexCD-OprM complex.

### 2.3 Overexpression of *mexCD* results in less available MexXY-OprM efflux pump

Having shown that DC interacts with NfxB inducing *mexCD* overexpression and that MexCD can form functional complexes with OprM, we next examined how increased *mexCD* expression might affect the activity of the aminoglycoside efflux pump MexXY-OprM. Our hypothesis was that an increased MexCD abundance may promote the formation of MexCD-OprM complexes, thereby sequestering OprM from MexXY and reducing the abundance of functional MexXY-OprM. This reduction would explain the increased and transient CS to tobramycin.

To test this hypothesis quantitatively, we built a simple mathematical assembly model including all three complexes: MexCD-OprJ, MexCD-OprM and MexXY-OprM (see Supplementary Note 2 for more details). We assumed that when *mexCD* is overexpressed, OprJ and OprM become limiting, resulting in competition particularly for OprM. This model (see Supplementary Note 2 for the derivation) leads to the following equilibrium concentration for MexXY-OprM:

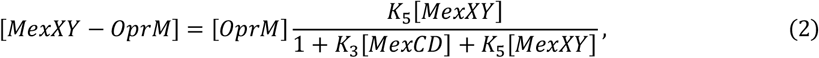

where *K*_*3*_ and *K*_*5*_ are equilibrium constants. The model predicts that as MexCD abundance increases, the concentration of MexXY-OprM decreases monotonically, whereas the concentrations of MexCD-OprJ and MexCD-OprM efflux pumps rise monotonically. Since the MexCD abundance itself increases with DC in a monotonic manner, both MexCD-containing efflux pumps inherit this dependence on DC.

### 2.4 Overexpression of *mexCD* results in transient resistance and collateral sensitivity to ciprofloxacin and tobramycin

We next examined how increased abundance of MexCD-OprJ and MexCD-OprM, together with reduced MexXY-OprM levels, may impact resistance to both ciprofloxacin and tobramycin, quantified as Minimal Inhibitory Concentrations (MICs). Using simple mathematical arguments (see Supplementary Note 3), we linked efflux pump abundance to MIC by assuming that bacterial growth rate decays as intracellular antibiotic concentration rises. Increased *mexCD* expression is expected to lower intracellular ciprofloxacin concentrations through overproduction of MexCD-OprJ and MexCD-OprM, while competition for OprM reduces MexXY-OprM levels and therefore raises intracellular tobramycin concentrations. From these assumptions we derived expressions relating *mexCD* expression levels to MICs for both antibiotics. Combining these with our expression for *mexCD* as a function of DC (equation (1)) yields:

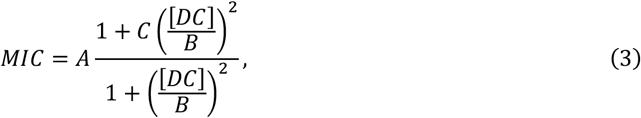

where *A, B*, and *C* are parameters to be determined (see Materials and Methods). The value of the parameter *C* sets the trend of the function: for ciprofloxacin *C*>1 and MIC increases with DC, for tobramycin *C*<1 and MIC decreases with DC (see Materials and Methods). Fitting this model using Bayesian estimation provides parameter posteriors and prediction intervals that closely align with our experimental MIC data (see Materials and Methods; Fig. 3), quantitatively validating the model prediction of a DC-dependent competition between MexCD and MexXY for the outer-membrane channel OprM.

**Figure 3.**
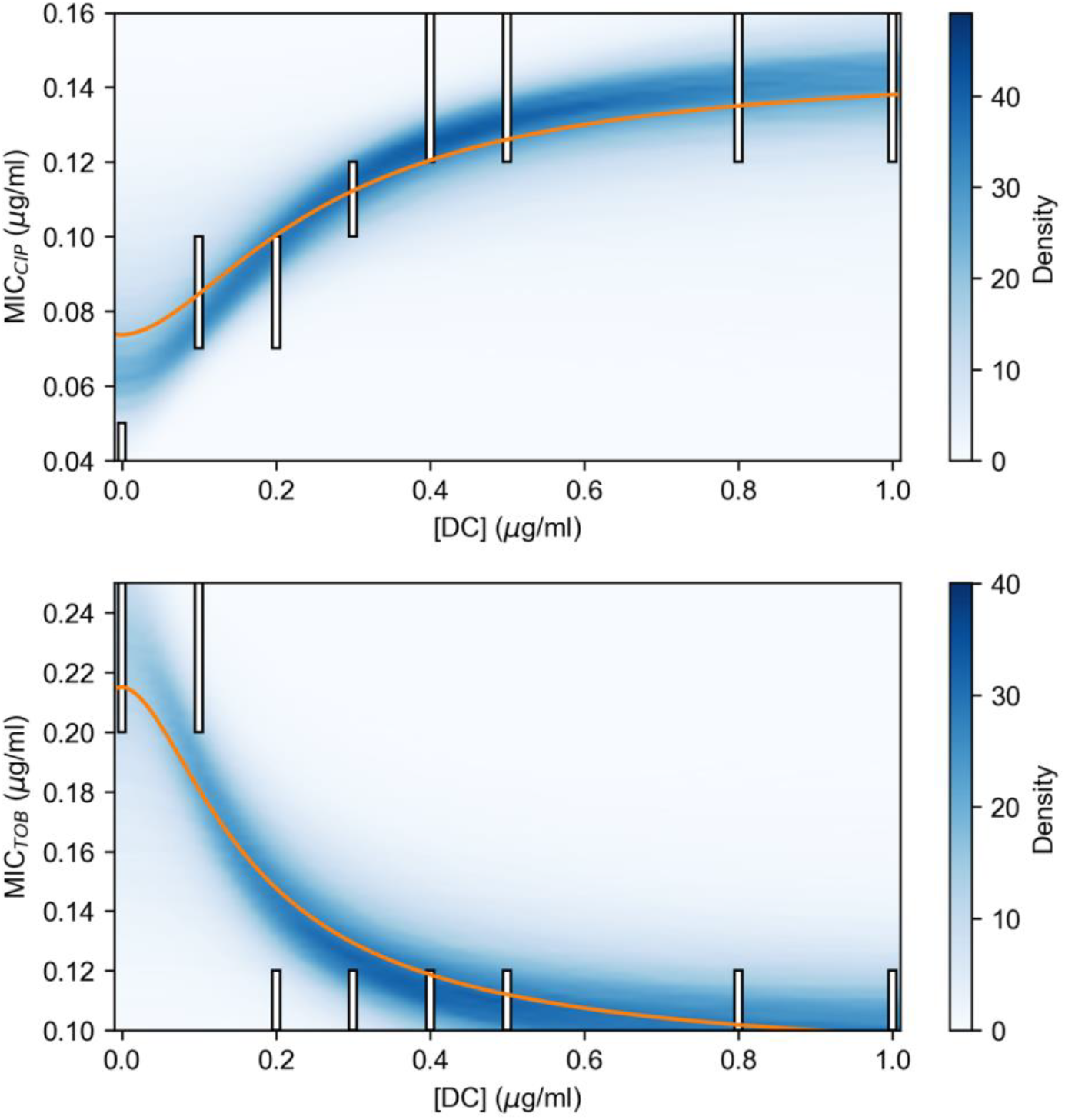
Minimal Inhibitory Concentrations (MICs) to ciprofloxacin (top) and tobramycin (bottom) as a function of DC. Bars indicate experimental MIC ranges obtained from checkerboard assays (see Materials and Methods). The blue shades represent the density of the posterior predictive distribution (i.e. the probability of a given MIC given a value of DC, according to the posterior probability distribution of the parameters *A, B, C* in equation (3)), calculated using Bayesian estimation (see Materials and Methods). Solid curves are the averages of the posterior predictive distributions for each value of DC. The fits quantitatively capture the expected trends: ciprofloxacin MIC increases with DC, whereas tobramycin MIC decreases with DC (see Materials and Methods).

### 2.5 Mutants lacking OprM do not show transient CS to aminoglycosides

Our model predicts that DC-dependent CS to tobramycin results from competition between MexCD and MexXY for the outer-membrane channel OprM. Accordingly, in the absence of OprM, DC-dependent CS to tobramycin should not occur. In contrast, in the absence of OprJ, CS to tobramycin should still be observed, while resistance to ciprofloxacin may be reduced. To experimentally test these predictions, we measured ciprofloxacin and tobramycin MICs for wild-type *P. aeruginosa* PA14 and isogenic mutants lacking either OprM or OprJ, in the presence or absence of DC (Fig. 4). MICs were determined using E-test strips, which are sensitive to modest shifts, and changes greater than two-fold increase or decrease were considered biologically relevant ^31,32.^

**Figure 4.**
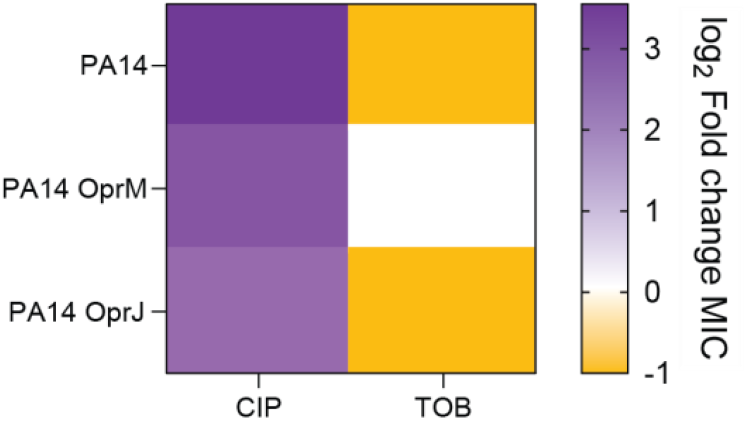
Outer-membrane channel-dependent effect of dequalinium chloride on ciprofloxacin and tobramycin susceptibility. Susceptibility to ciprofloxacin and tobramycin was assessed in the *P. aeruginosa* PA14 wild-type strain and in PA14OprM and PA14OprJ mutants, in the presence or absence of dequalinium chloride (DC). Color intensity represents the log-transformed fold change in MIC in the presence of DC relative to the MIC of the respective parental strain without DC. Changes in MIC greater than or equal to a 2-fold increase or decrease were considered biologically relevant and used to classify populations as “resistant” (purple) or “susceptible” (yellow), as previously described ^19^. MIC values (µg/mL) are provided in Supplementary Table 2. CIP, ciprofloxacin; TOB, tobramycin.

In line with our previous works ^27^, DC exposure increased ciprofloxacin resistance and induced CS to tobramycin in wild-type PA14. In the OprJ mutant, DC still raised ciprofloxacin MICs and elicited CS to tobramycin (Fig. 4). This result is consistent with the fact that MexCD, typically paired with OprJ, can also interact with alternative outer-membrane partners, such as OprM, to mediate ciprofloxacin efflux, causing resistance levels that are lower than that in the presence of its canonical outer-membrane protein OprJ. These findings support our model prediction that in the absence of OprJ, MexCD relies on OprM. This dependence increases competition with MexXY for OprM, thereby reducing the abundance of functional MexXY-OprM complexes and increasing susceptibility to tobramycin. In contrast, the OprM mutant, which lacks the outer-membrane protein required for MexXY function, showed increased ciprofloxacin resistance upon DC exposure but did not exhibit CS to tobramycin (Fig. 4). This observation is also consistent with our model prediction: in the absence of OprM, MexCD cannot use this outer-membrane channel even when DC causes a significant demand for the MexCD efflux pump, and, therefore, tobramycin resistance does not shift. As expected, if both OprM and OprJ contribute to MexCD activity when efflux demand is high, this mutant also exhibited a reduced increase of ciprofloxacin MIC compared to wild-type PA14 after DC exposure. Together, these results indicate that competition for outer-membrane components is a constitutive feature of *P. aeruginosa* physiology that generates important trade-offs when high levels of MexCD efflux activity are required (Conceptual Fig. 5).

**Figure 5.**
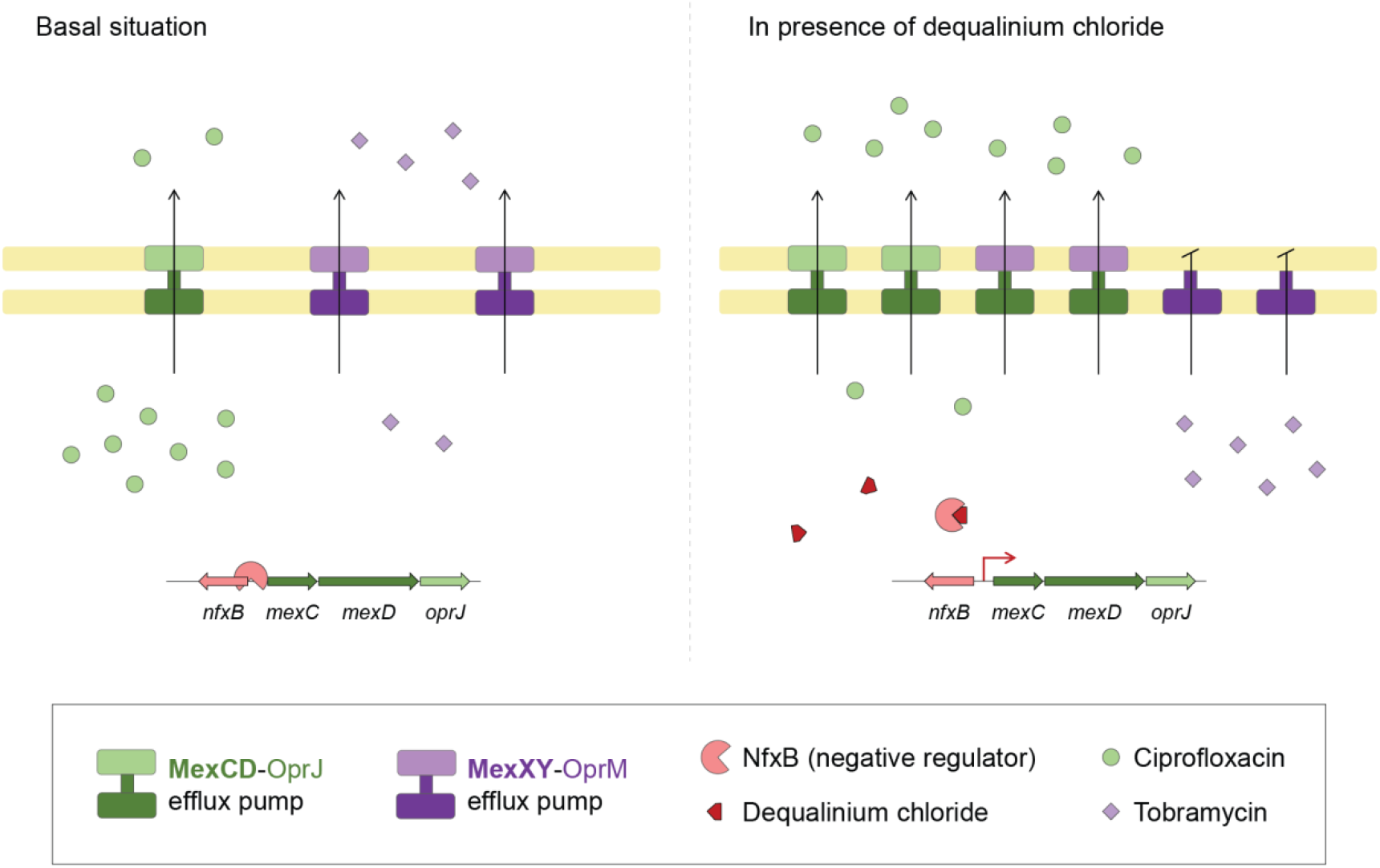
Conceptual representation of the molecular mechanism responsible for transient collateral sensitivity to tobramycin induced by dequalinium chloride on *P. aeruginosa*. Dequalinium chloride (DC) binds to NfxB, relieving its repression of *mexCD-oprJ* expression. This leads to increased formation of the MexCD efflux pump, which is assembled with either OprJ or OprM porins, thereby enhancing ciprofloxacin efflux. Overproduction of MexCD sequesters OprM from the MexXY efflux pump, reducing the assembly of MexXY-OprM in the membrane and compromising tobramycin extrusion.

## 3 Discussion

The global health challenge posed by AMR demands new strategies to enhance antibiotic efficacy ^38^. CS is an evolutionary trade-off associated with AMR and offers a promising framework for designing rational treatment regimens. However, historical contingency complicates the implementation of CS-based strategies. For example, an analysis of nearly 450,000 susceptibility tests collected over four years from hospitals in Pittsburgh found that variation in CS patterns undermines the effectiveness of therapies based on alternating antibiotics ^39^.

In previous work, we addressed this limitation by deliberately inducing transient CS to tobramycin via overproduction of the MexCD-OprJ efflux pump. The rationale was straightforward: rather than relying on ciprofloxacin to select loss-of function mutations in *nfxB*, and potentially other unintended targets, we employed DC to transiently inhibit NfxB activity ^27^. This strategy offers several advantages over approaches based on selecting stable resistance mutations. It bypasses historical contingency by directly targeting a specific AMR mechanism and avoids the risk of selecting antibiotic-resistant mutants if treatment fails. Our findings also expanded the concept of CS, a term coined in the 1950s as increased susceptibility to an antibiotic following acquisition of resistance to another ^40^, including cases where the increased susceptibility to the second antibiotic is not stable.

Overexpression of *mexCD-oprJ* can arise either from the acquisition of ciprofloxacin-selected loss-of-function mutations in its local repressor gene *nfxB* ^17,37,41^ or from inducer compounds that interfere with the NfxB repressive activity ^27^. The antiseptic and disinfectant DC is a potent inducer of *mexCD-oprJ* expression ^32^ that binds and inhibits the TetR-family regulator RamR ^35^. Here we provide structural and quantitative evidence consistent with DC binding to NfxB and inhibiting its regulatory function. Docking analyses suggest that it could disrupt protein-DNA interactions, and predicted binding affinities indicate that DC binds NfxB with higher affinity than RamR. Together with our mechanistic mathematical model of DC-NfxB binding and its impact on *mexCD-oprJ* expression, which accurately captures the experimental data, these results support a mechanism in which DC induces transient CS to tobramycin by inhibiting NfxB-mediated repression.

Structural simulations showed that MexCD can assemble with the outer-membrane channel OprM, supporting the possibility of competition for this component with the MexXY-OprM efflux pump. Interpreting efflux-pump simulations requires caution because we cannot explicitly model the cell wall, as current understanding of its detailed architecture remains insufficient to build accurate atomic models ^42^. As an approximation, our simulations were performed in saline solution omitting many *in vivo* constraints that likely stabilize the assemblies. Paradoxically, that makes our observations more compelling: if the complexes remain bound and stable without those constraints, their stability would likely be even higher in the presence of the cell wall. Over simulated timescales up to 100 ns, our results support that the MexCD-OprJ and MexCD-OprM interactions are comparable in stability to the reference MexAB-OprM complex.

Our mechanistic model of efflux-pump assembly predicts that DC-enhanced *mexCD* expression reduces the availability of MexXY-OprM complexes through competition for the shared outer-membrane channel. After relating efflux pumps levels to resistance, our mathematical model quantitatively reproduces the observed MIC shifts for both ciprofloxacin and tobramycin. Using a minimal set of assumptions, the model delivers accurate MIC predictions, supporting the quantitative validity of our mechanistic hypothesis. Importantly, the framework is parsimonious yet extensible, providing a transparent basis for exploring alternative regulatory scenarios. It also yields clear, testable predictions: for example, lower OprM availability should eliminate the effect (increased susceptibility) of DC, whereas lower OprJ should not cause changes with respect to the wild type. We tested these two predictions using two mutants lacking OprM or OprJ in the wild-type *P. aeruginosa* PA14 strain, providing strong experimental support for the hypothesis that CS to tobramycin results from competition between MexCD and MexXY efflux pumps for the shared outer-membrane protein OprM (Fig. 5).

Overall, our results provide a mechanistic explanation for DC-associated transient CS to tobramycin and suggest new therapeutic opportunities. In addition to targeting transcriptional regulators such as NfxB, compounds that modulate outer-membrane channel availability may provide alternative strategies for inducing CS. More broadly, identifying inducible CS mechanisms and elucidating their molecular basis could help improve antibiotic efficacy and the treatment of bacterial infections. Indeed, transient CS may be particularly attractive therapeutically because it bypasses historical contingency and minimizes the risk of selecting antibiotic-resistant mutants. These findings provide a proof of concept that mechanistic understanding of resistance trade-offs can inform new approaches to antimicrobial therapy.

Finally, the ability of outer-membrane components to interact with multiple efflux systems likely extends beyond *P. aeruginosa*, suggesting that similar competition-driven trade-offs may occur in other bacterial pathogens. Indeed, the amplification of the AcrAB efflux operon in *E. coli* was recently linked to CS to gentamicin ^43^, although the underlying mechanism remains elusive. Our results suggest that this phenomenon may stem from competition with the AcrAD efflux pump for the shared outer-membrane component TolC ^44^, an hypothesis that we will explore in future work. Consequently, our findings provide a general mechanistic framework for understanding and potentially exploiting inducible CS mediated by multidrug-resistance efflux pumps.

## 4 Materials and Methods

### 4.1 Bacterial strains and growth experiments

*P. aeruginosa* PA14 wild-type strain and mutants lacking OprM or OprJ outer-membrane proteins (PA14 Tn OprM and PA14 Tn OprJ), from a non-redundant PA14 transposon insertion library ^45^, were grown in glass tubes in Lysogeny Broth (LB; Lenox, Pronadisa) at 37 °C with shaking at 250 rpm. MICs of ciprofloxacin and tobramycin were determined at 37 °C, in Mueller Hinton (MH) agar, using E-test strips (MIC Test Strip, Liofilchem®) in the presence or absence of 10 μg/mL of DC, as previously described ^27^. Cells were pre-grown in LB supplemented or not with 10 μg/mL of DC prior to MIC determination. Checkerboard broth microdilution assays were performed in 96-U-well plates using MH medium containing serial dilutions of DC (0.1 to 1 μg/mL), tobramycin (0.031 to 0.25 μg/mL) or ciprofloxacin (0.013 to 0.125 μg/mL). Each well contained 90 µL medium and was inoculated with 10 μL of bacterial suspensions to a final OD_600nm_ of 0.01. Plates were incubated at 37 °C for 48 h without shaking.

### 4.2 Modeling of NfxB binding to dequalinium chloride and estimation of affinities

The structure of DC was retrieved from the ZINC database (id:194130902), and MOPAC was used to optimize its fine structure and calculate its charges. Models of *P. aeruginosa* NfxB were built using AlphaFold-2, ESMFold, OmegaFold, I-TASSER and retrieved from SwissModel ^46^ and ModBase, preprocessed with UCSF-Chimera “DockPrep” tool, docked against DC using DOCK6, FireDock and AutoDock Vina, and further refined with a minimization step using the AMBER force field ^47^ in UCSF-Chimera, prior to manual inspection and selection of representative poses. The predicted structure for NfxB is close to that of the published structures of RamR (PDB:3VW0, RMSD 1.11Å).

The structures of RamR bound to berberine, chenodeoxycolic acid, dequalinium and rhodamine 6G were retrieved from RCSB-PDB ^48^ (entries 3VVZ, 3VW0 and 6IE9). The affinity of RamR for berberine, chenodeoxycolic acid, dequalinium and rhodamine 6G, and of the poses selected after docking NfxB and dequalinium was calculated using AutoDock Vina, NNScore ^49^ and DLScore, and the measures averaged for estimation.

### 4.3 Dynamic Modeling of Mex-Opr complexes

Initial structures for *P. aeruginosa* MexAB-OprM complex (entries 6IOK, 6IOL, 6TA5 6TA6), MexA (1T5E, 1VF7, 2V4D), MexB (2V50, 3W9I, 3W9Jm 6IIA), OprM (1WP1, 3D5K, 4Y1K, 6IOK, 6IOL, 6TA5, 6TA6, 6ZRE, 7AKZ), OprJ (5AZS), OprN (5AZ0, SAZP) and MexY (9E9F) were retrieved from RCSB-PDB.

Monomers lacking a known structure were first modeled: the structures of MexC, MexD and MexX were predicted using AlphaFold, ESMFold and Modeller. Selection of the initial structures and models was based on preservation of features (relative orientations and interactions), and on their amenability to assembly (preservation of joint features like the central pore tunnel, reduction of steric conflicts, relative positions among subunits, conservation of contacts, similarity to known reference structures and absence of physical artifacts). In each case, all the available structures were tried for assemblage in complexes and the most suitable structure was selected. Complexes were incrementally built, first as homo-hexamers or homo-trimers (as required), then as MexCD or MexXY complexes and finally as OprM or OprJ with MexC, MexX or MexCD complexes. Final assemblies were minimized in order to allow the components to relax in their mutual presence prior to molecular dynamics simulation. The complete assemblies were produced superposing the structures over the known structure of MexAB-OprM (Figure 3), then separating homo-multimers to avoid steric clashes and finally minimizing the structure *in vacuo* to allow it to refit itself.

To evaluate the stability of the complexes built, we subjected them to long molecular dynamics simulations as follows: initial structure and complex minimizations were done with the CHARMM force field ^50^ in GROMACS ^51^ using a complex imbibed in 150mM saline solution. Fully atomistic simulations were also performed using CHARMM in GROMACS using the complex imbibed in 150mM saline solution (approx. 15 million atoms), with a minimization (up to 10000 steps), equilibration in the NVT (200ps in 2 fs steps) and NPT (200 ps in 2 fs steps) ensembles, and a molecular dynamics (MD) production run of 5 ns in 2 fs steps, at 1 bar and 310 °K. Once fully atomistic models were deemed suitable, to achieve longer simulation times, coarse-grained simulations were performed using the Martini force field ^52^ in GROMACS, with the complex imbibed in 150mM saline solution, using a cubic box 200 nm on each side (approx. 0.75-0.8 million particles), and applying one or more (as needed) minimizations (5000 steps), an equilibration phase (10000 steps of 10 fs) and a production MD phase (5000000 steps of 20 fs, 100 ns). After coarse-grained simulation, fully atomistic structures were reconstructed using Pulchra ^53^.

### 4.4. Bayesian estimation

We used Bayesian inference to estimate the posterior distribution of the parameters in equations (1) and (3) in relation to our laboratory data. In order to do this, we employed the NUTS algorithm as implemented in the Python library PyMC ^54^. For Figure 3, we took the MIC to be the average between the minimal concentration of antibiotic that inhibited growth and the maximal concentration that allowed growth, understanding that the actual MIC could be anywhere between the two concentrations. For instance, if a concentration 0.125 μg/mL allowed growth but the next concentration measured (0.25 μg/mL) did not, we take the MIC to be 0.1875 μg/mL.

## 4.5. Data availability

Scripts used to drive the molecular structure simulations are available in folder MolMod of github.com/jrvalverde/MolModTools. Codes for reproducing the inference and for generating the associated figures (1B and 3) can be found at github.com/pablocatalan/inducers. Data needed to evaluate the conclusions of this work are present in the manuscript and Supplementary Materials.

## 4.6. Acknowledgments

Work was supported by grants PID2022-142185NB-C22 to PC, PID2022-142185NB-C21 to SA, PID2023-149913OA-I00 to SHA, funded by the Spanish Ministerio de Ciencia e Innovación (MCIN/AEI/10.13039/501100011033) and by ERDF/EU ‘A way of making Europe’. PL is the recipient of an ERS/EU RESPIRE4 Marie Skłodowska-Curie Postdoctoral Research Fellowship (Ref. nr: R4202305-01047). The MICIU/AEI/10.13039/501100011033 is acknowledged for supporting the National Center for Biotechnology through the “Severo Ochoa” grant CEX2023-001386-S.

## Author contributions

P.C., S.A., J.L.M., and S.H.-A. conceived the study and designed the research. P.C., S.A., and B.P.-E. developed and performed mathematical modelling. J.R.V. and I.K. performed the structural simulations. S.H.-A. and P.L. performed the experiments. S.H.-A., P.L., P.C., S.A., J.R.V., and J.L.M. wrote the manuscript, with contributions and final approval from all authors.

## REFERENCES

1 Catalan, P. et al. Seeking patterns of antibiotic resistance in ATLAS, an open, raw MIC database with patient metadata. Nature communications 13, 2917, doi:10.1038/s41467-022-30635-7 (2022).

2 Collaborators, G. B. D. A. R. Global mortality associated with 33 bacterial pathogens in 2019: a systematic analysis for the Global Burden of Disease Study 2019. Lancet 400, 2221–2248, doi:10.1016/S0140-6736(22)02185-7 (2022).

3 Antimicrobial Resistance, C. Global burden of bacterial antimicrobial resistance in 2019: a systematic analysis. Lancet 399, 629–655, doi:10.1016/S0140-6736(21)02724-0 (2022).

4 Martinez-Solano, L., Macia, M. D., Fajardo, A., Oliver, A. & Martinez, J. L. Chronic *Pseudomonas aeruginosa* infection in chronic obstructive pulmonary disease. Clinical Infectious Diseases 47, 1526–1533 (2008).

5 Nickerson, R., Thornton, C. S., Johnston, B., Lee, A. H. Y. & Cheng, Z. Pseudomonas aeruginosa in chronic lung disease: untangling the dysregulated host immune response. Frontiers in immunology 15, 1405376, doi:10.3389/fimmu.2024.1405376 (2024).

6 Olivares, J. et al. The intrinsic resistome of bacterial pathogens. Frontiers in microbiology 4, 103, doi:10.3389/fmicb.2013.00103 (2013).

7 Bellido, F., Martin, N. L., Siehnel, R. J. & Hancock, R. E. Reevaluation, using intact cells, of the exclusion limit and role of porin OprF in Pseudomonas aeruginosa outer membrane permeability. J Bacteriol 174, 5196–5203, doi:10.1128/jb.174.16.5196-5203.1992 (1992).

8 Lopez-Causape, C., Cabot, G., Del Barrio-Tofino, E. & Oliver, A. The Versatile Mutational Resistome of Pseudomonas aeruginosa. Frontiers in microbiology 9, 685, doi:10.3389/fmicb.2018.00685 (2018).

9 Lopez-Causape, C. et al. Evolution of the Pseudomonas aeruginosa mutational resistome in an international Cystic Fibrosis clone. Sci Rep 7, 5555, doi:10.1038/s41598-017-05621-5 (2017).

10 Baym, M., Stone, L. K. & Kishony, R. Multidrug evolutionary strategies to reverse antibiotic resistance. Science 351, aad3292, doi:10.1126/science.aad3292 (2016).

11 Herencias, C. et al. beta-lactamase expression induces collateral sensitivity in Escherichia coli. Nature communications 15, 4731, doi:10.1038/s41467-024-49122-2 (2024).

12 Herencias, C. et al. Collateral sensitivity associated with antibiotic resistance plasmids. eLife 10, doi:10.7554/eLife.65130 (2021).

13 Pal, C., Papp, B. & Lazar, V. Collateral sensitivity of antibiotic-resistant microbes. Trends in microbiology 23, 401–407, doi:10.1016/j.tim.2015.02.009 (2015).

14 Munck, C., Gumpert, H. K., Wallin, A. I., Wang, H. H. & Sommer, M. O. Prediction of resistance development against drug combinations by collateral responses to component drugs. Science translational medicine 6, 262ra156, doi:10.1126/scitranslmed.3009940 (2014).

15 Barbosa, C., Beardmore, R., Schulenburg, H. & Jansen, G. Antibiotic combination efficacy (ACE) networks for a Pseudomonas aeruginosa model. PLoS biology 16, e2004356, doi:10.1371/journal.pbio.2004356 (2018).

16 Imamovic, L. & Sommer, M. O. Use of collateral sensitivity networks to design drug cycling protocols that avoid resistance development. Science translational medicine 5, 204ra132, doi:10.1126/scitranslmed.3006609 (2013).

17 Imamovic, L. et al. Drug-Driven Phenotypic Convergence Supports Rational Treatment Strategies of Chronic Infections. Cell 172, 121–134 e114, doi:10.1016/j.cell.2017.12.012 (2018).

18 Kim, S., Lieberman, T. D. & Kishony, R. Alternating antibiotic treatments constrain evolutionary paths to multidrug resistance. Proceedings of the National Academy of Sciences of the United States of America 111, 14494–14499, doi:10.1073/pnas.1409800111 (2014).

19 Hernando-Amado, S., Laborda, P., Valverde, J. R. & Martinez, J. L. Mutational background influences P. aeruginosa ciprofloxacin resistance evolution but preserves collateral sensitivity robustness. Proceedings of the National Academy of Sciences of the United States of America 119, e2109370119, doi:10.1073/pnas.2109370119 (2022).

20 Hernando-Amado, S. et al. Rapid Phenotypic Convergence towards Collateral Sensitivity in Clinical Isolates of Pseudomonas aeruginosa Presenting Different Genomic Backgrounds. Microbiology spectrum, e0227622, doi:10.1128/spectrum.02276-22 (2022).

21 Hernando-Amado, S., Sanz-Garcia, F. & Martinez, J. L. Rapid and robust evolution of collateral sensitivity in Pseudomonas aeruginosa antibiotic-resistant mutants. Science advances 6, eaba5493, doi:10.1126/sciadv.aba5493 (2020).

22 Podnecky, N. L. et al. Conserved collateral antibiotic susceptibility networks in diverse clinical strains of Escherichia coli. Nature communications 9, 3673, doi:10.1038/s41467-018-06143-y (2018).

23 Barbosa, C. et al. Alternative Evolutionary Paths to Bacterial Antibiotic Resistance Cause Distinct Collateral Effects. Molecular biology and evolution 34, 2229–2244, doi:10.1093/molbev/msx158 (2017).

24 Lazar, V. et al. Bacterial evolution of antibiotic hypersensitivity. Molecular systems biology 9, 700, doi:10.1038/msb.2013.57 (2013).

25 Lazar, V. et al. Genome-wide analysis captures the determinants of the antibiotic cross-resistance interaction network. Nature communications 5, 4352, doi:10.1038/ncomms5352 (2014).

26 Armstrong, D. et al. Evidence for spread of a clonal strain of Pseudomonas aeruginosa among cystic fibrosis clinics. Journal of clinical microbiology 41, 2266–2267, doi:10.1128/JCM.41.5.2266-2267.2003 (2003).

27 Hernando-Amado, S., Laborda, P. & Martinez, J. L. Tackling antibiotic resistance by inducing transient and robust collateral sensitivity. Nature communications 14, 1723, doi:10.1038/s41467-023-37357-4 (2023).

28 Jeannot, K. et al. Resistance and virulence of Pseudomonas aeruginosa clinical strains overproducing the MexCD-OprJ efflux pump. Antimicrobial agents and chemotherapy 52, 2455–2462, doi:10.1128/AAC.01107-07 (2008).

29 Morita, Y., Tomida, J. & Kawamura, Y. MexXY multidrug efflux system of Pseudomonas aeruginosa. Frontiers in microbiology 3, 408, doi:10.3389/fmicb.2012.00408 (2012).

30 Adewoye, L., Sutherland, A., Srikumar, R. & Poole, K. The mexR repressor of the mexAB-oprM multidrug efflux operon in Pseudomonas aeruginosa: characterization of mutations compromising activity. J Bacteriol 184, 4308–4312, doi:10.1128/JB.184.15.4308-4312.2002 (2002).

31 Gotoh, N. et al. Functional replacement of OprJ by OprM in the MexCD-OprJ multidrug efflux system of Pseudomonas aeruginosa. FEMS microbiology letters 165, 21–27, doi:10.1111/j.1574-6968.1998.tb13122.x (1998).

32 Laborda, P., Alcalde-Rico, M., Blanco, P., Martínez, J. L. & Hernando-Amado, S. Novel Inducers of the Expression of Multidrug Efflux Pumps That Trigger Pseudomonas aeruginosa Transient Antibiotic Resistance. Antimicrobial agents and chemotherapy 63, doi:10.1128/aac.01095-19 (2019).

33 Monti, M. R., Morero, N. R., Miguel, V. & Argarana, C. E. nfxB as a novel target for analysis of mutation spectra in Pseudomonas aeruginosa. PLoS One 8, e66236, doi:10.1371/journal.pone.0066236 (2013).

34 Purssell, A. & Poole, K. Functional characterization of the NfxB repressor of the mexCD-oprJ multidrug efflux operon of Pseudomonas aeruginosa. Microbiology 159, 2058–2073, doi:10.1099/mic.0.069286-0 (2013).

35 Yamasaki, S. et al. The crystal structure of multidrug-resistance regulator RamR with multiple drugs. Nature communications 4, 2078, doi:10.1038/ncomms3078 (2013).

36 Bienert, S. et al. The SWISS-MODEL Repository-new features and functionality. Nucleic acids research 45, D313–D319, doi:10.1093/nar/gkw1132 (2017).

37 Masuda, N. et al. Substrate specificities of MexAB-OprM, MexCD-OprJ, and MexXY-oprM efflux pumps in Pseudomonas aeruginosa. Antimicrobial agents and chemotherapy 44, 3322–3327 (2000).

38 Laxminarayan, R. Antibiotic effectiveness: balancing conservation against innovation. Science 345, 1299–1301, doi:10.1126/science.1254163 (2014).

39 Beckley, A. M. & Wright, E. S. Identification of antibiotic pairs that evade concurrent resistance via a retrospective analysis of antimicrobial susceptibility test results. The Lancet. Microbe 2, e545–e554, doi:10.1016/s2666-5247(21)00118-x (2021).

40 Szybalski, W. & Bryson, V. Genetic studies on microbial cross resistance to toxic agents. I. Cross resistance of Escherichia coli to fifteen antibiotics. J Bacteriol 64, 489–499 (1952).

41 Poole, K. et al. Overexpression of the *mexC-mexD-oprJ* efflux operon in *nfxB*-type multidrug-resistant strains of *Pseudomonas aeruginosa*. Mol Microbiol 21, 713–724 (1996).

42 Rohde, M. The Gram-Positive Bacterial Cell Wall. Microbiology spectrum 7, doi:10.1128/microbiolspec.GPP3-0044-2018 (2019).

43 Garoff, L. et al. Population Bottlenecks Strongly Influence the Evolutionary Trajectory to Fluoroquinolone Resistance in Escherichia coli. Molecular biology and evolution 37, 1637–1646, doi:10.1093/molbev/msaa032 (2020).

44 Garneau-Tsodikova, S. & Labby, K. J. Mechanisms of Resistance to Aminoglycoside Antibiotics: Overview and Perspectives. MedChemComm 7, 11–27, doi:10.1039/C5MD00344J (2016).

45 Liberati, N. T. et al. An ordered, nonredundant library of Pseudomonas aeruginosa strain PA14 transposon insertion mutants. Proceedings of the National Academy of Sciences of the United States of America 103, 2833–2838, doi:10.1073/pnas.0511100103 (2006).

46 Kiefer, F., Arnold, K., Kunzli, M., Bordoli, L. & Schwede, T. The SWISS-MODEL Repository and associated resources. Nucleic acids research 37, D387–392, doi:10.1093/nar/gkn750 (2009).

47 Wang, J., Wolf, R. M., Caldwell, J. W., Kollman, P. A. & Case, D. A. Development and testing of a general amber force field. Journal of computational chemistry 25, 1157–1174, doi:10.1002/jcc.20035 (2004).

48 Rose, P. W. et al. The RCSB protein data bank: integrative view of protein, gene and 3D structural information. Nucleic acids research 45, D271–D281, doi:10.1093/nar/gkw1000 (2017).

49 Durrant, J. D. & McCammon, J. A. NNScore 2.0: a neural-network receptor-ligand scoring function. Journal of chemical information and modeling 51, 2897–2903, doi:10.1021/ci2003889 (2011).

50 Zhu, X., Lopes, P. E. & Mackerell, A. D., Jr. Recent Developments and Applications of the CHARMM force fields. Wiley interdisciplinary reviews. Computational molecular science 2, 167–185, doi:10.1002/wcms.74 (2012).

51 Abraham, M. J. et al. GROMACS: High performance molecular simulations through multi-level parallelism from laptops to supercomputers. SoftwareX 1-2, 19–25 (2015).

52 Periole, X. & Marrink, S. J. The Martini coarse-grained force field. Methods in molecular biology 924, 533–565, doi:10.1007/978-1-62703-017-5_20 (2013).

53 Rotkiewicz, P. & Skolnick, J. Fast procedure for reconstruction of full-atom protein models from reduced representations. Journal of computational chemistry 29, 1460–1465, doi:10.1002/jcc.20906 (2008).

54 Abril-Pla, O. et al. PyMC: a modern, and comprehensive probabilistic programming framework in Python. PeerJ. Computer science 9, e1516, doi:10.7717/peerj-cs.1516 (2023).

