## Supplementary Material for "Exploiting outer-membrane protein promiscuity to induce transient collateral sensitivity via efflux pump competition"

[1]Grupo Interdisciplinar de Sistemas Complejos (GISC), Madrid, Spain.

[2]Departamento de Matemáticas, Universidad Carlos III de Madrid, 28911 Leganés, Spain.

[3]Centro Nacional de Biotecnologia (CNB), CSIC, 28049 Madrid, Spain.

[4]Dpto. de Matemáticas e Informática (DMIAICN), ETSI Caminos, Canales y Puertos, Universidad Politécnica de Madrid, C/ Prof. Aranguren 3, 28040 Madrid, Spain.

[5]Department of Clinical Microbiology 9301, Rigshospitalet, 2100 Copenhagen, Denmark.

∗ corresponding authors:

PC:

SHA:

**Supplementary Note 1: expression of *mexCD* as a function of DC**

The known relationships between DNA promoter ($G$), NfxB ($N$, in its tetrameric form) and DC ($DC$) can be written, in a simplified way, as the following chemical reactions:

$$\begin{aligned} G + N \quad & \underset{k_{-1}}{\overset{k_1}{\rightleftharpoons}} C_1 \\ 2DC + N \quad & \underset{k_{-2}}{\overset{k_2}{\rightleftharpoons}} C_2 \\ DC_{out} \quad & \underset{\phi_{out}}{\overset{\phi_{in}}{\rightleftharpoons}} DC \end{aligned} \tag{4}$$

where $k_i$ are reaction rates, $DC_{out}$ is extracellular DC, $C_1$ is the complex formed by NfxB and the *mexCD* promoter region, and $C_2$ is the DC-NfxB complex. Using mass action kinetics, and assuming the extracellular DC pool is very large, we can write these reactions as a set of ordinary differential equations:

$$\begin{aligned} \frac{d[G]}{dt} &= -k_1[G][N] + k_{-1}[C_1] \\ \frac{d[N]}{dt} &= -k_1[G][N] + k_{-1}[C_1] - k_2[DC]^2[N] + k_{-2}[C_2] \\ \frac{d[DC]}{dt} &= \phi_{in}[DC_{out}] - \phi_{out}[DC] - 2k_2[DC]^2[N] + 2k_{-2}[C_2] \\ \frac{d[C_1]}{dt} &= k_1[G][N] - k_{-1}[C_1] \\ \frac{d[C_2]}{dt} &= k_2[DC]^2[N] - k_{-2}[C_2] \end{aligned} \tag{5}$$

where $[\cdot]$ represents concentration, $k_i$ are the reaction rates and $[DC_{out}]$ is taken as a constant. Note that $[G] + [C_1] = [G_{tot}]$ and $[N] + [C_2] = [N_{tot}]$ are constants and represent the total (bound or unbound) concentration of promoter and NfxB in the cell, respectively.

Assuming steady state (all derivatives equal to zero), we obtain from the equations for $[G]$ and $[C_1]$ an expression for the stationary concentration $[G]_*$ as a function of $[N]_*$:

$$[G]_* = \frac{[G_{tot}]}{1 + \frac{k_1}{k_{-1}}[N]_*}. \qquad (6)$$

Similarly, from the equations for $[N]$ and $[C_2]$ we obtain an expression for the stationary concentration $[N]_*$ as a function of $[DC]_*$:

$$[N]_* = \frac{[N_{tot}]}{1 + \frac{k_2}{k_{-2}}[DC]_*^2}. \qquad (7)$$

We can substitute equation (7) into equation (6) to obtain

$$[G]_* = [G_{tot}] \frac{1 + \frac{k_2}{k_{-2}}[DC]_*^2}{1 + \frac{k_1}{k_{-1}}[N_{tot}] + \frac{k_2}{k_{-2}}[DC]_*^2}. \qquad (8)$$

Finally, from the equation for $[DC]$ we obtain

$$[DC]_* = \frac{\phi_{in}}{\phi_{out}} DC_{out}. \qquad (9)$$

Since *mexCD* ($MexCD$) expression depends linearly on $[G]_*$, we can use all the previous information to deduce a Hill relationship between $MexCD$ and $DC_{out}$:

$$[MexCD] = a\frac{1 + \left(\frac{[DC_{out}]}{b}\right)^2}{c + \left(\frac{[DC_{out}]}{b}\right)^2}, \tag{10}$$

Where $a$, $b$, $c$ are positive unknown parameters. This is equation (1) in the main text. We can determine the unknown parameters fitting equation (10) to previous *mexCD* expression data from our laboratory [1]. The fit obtained (Figure 1B) aligns well with the experimental data and supports the proposed model.

**Supplementary Note 2: relation between MexCD and MexXY efflux pumps**

We model the association and dissociation of efflux pumps, representing a promiscuous system where all possible types of complexes of MexCD and MexXY, on one hand, and OprM and OprJ, on the other hand, can form, resulting in the following system of ordinary differential equations:

$$\begin{aligned}
\frac{dCD}{dt} &= -k_1 CD\, J + k_2 P_1 - k_3 CD\, M + k_4 P_2 \\
\frac{dJ}{dt} &= -k_1 CD\, J + k_2 P_1 - k_7 XY\, J + k_8 P_4 \\
\frac{dXY}{dt} &= -k_5 XY\, M + k_6 P_3 - k_7 XY\, J + k_8 P_4 \\
\frac{dM}{dt} &= -k_3 CD\, M + k_4 P_2 - k_5 XY\, M + k_6 P_3 \\
\frac{dP_1}{dt} &= k_1 CD\, J - k_2 P_1 \\
\frac{dP_2}{dt} &= k_3 CD\, M - k_4 P_2 \\
\frac{dP_3}{dt} &= k_5 XY\, M - k_6 P_3 \\
\frac{dP_4}{dt} &= k_7 XY\, J - k_8 P_4.
\end{aligned} \tag{11}$$

where $CD$ is the concentration of free MexCD complex, $J$ is the concentration of free OprJ; $XY$ is the concentration of free MexXY complex; $M$ is the concentration of free OprM; $P_1$ is the concentration of the efflux pump MexCD-OprJ; $P_2$ is the concentration of the efflux pump MexCD-OprM; $P_3$ is the concentration of the efflux pump MexXY-OprM; $P_4$ is the concentration of the efflux pump MexXY-OprJ; $k_1$, $k_3$, $k_5$ and $k_7$ are association constants, whereas $k_2$, $k_4$, $k_6$ and $k_8$ are dissociation constants. As seen in Figure 2, the stoichiometry to form the efflux pumps is 1. In the Main Text we do not consider the formation of a MexXY-OprJ efflux pump: this is recovered taking $k_7 = 0$ in equations (11). Here we start discussing the general case assuming $k_7 \geq 0$.

At steady-state, we set $\frac{dCD}{dt} = \frac{dJ}{dt} = \frac{dXY}{dt} = \frac{dM}{dt} = \frac{dP_1}{dt} = \frac{dP_2}{dt} = \frac{dP_3}{dt} = \frac{dP_4}{dt} = 0$. From the steady-state conditions:

$$\begin{aligned} k_1 CD\ J &= k_2 P_1 \Rightarrow P_1 = \frac{k_1}{k_2} CD\ J \\ k_3 CD\ M &= k_4 P_2 \Rightarrow P_2 = \frac{k_3}{k_4} CD\ M \\ k_5 XY\ M &= k_6 P_3 \Rightarrow P_3 = \frac{k_5}{k_6} XY\ M \\ k_7 XY\ J &= k_8 P_4 \Rightarrow P_4 = \frac{k_7}{k_8} XY\ J. \end{aligned} \tag{12}$$

The total quantity of *CD* (or *XY*) is in free form or forming part of efflux pumps $P_1$ and $P_2$ ($P_3$ and $P_4$), so we can define variables for the total concentrations:

$$\begin{aligned} CD_0 &\equiv CD + P_1 + P_2 \\ XY_0 &\equiv XY + P_3 + P_4. \end{aligned} \tag{13}$$

Using this, and defining the equilibrium constants:

$$K_1 \equiv \frac{k_1}{k_2}, \quad K_3 \equiv \frac{k_3}{k_4}, \quad K_5 \equiv \frac{k_5}{k_6}, \quad K_7 \equiv \frac{k_7}{k_8}, \tag{14}$$

we can write:

$$
\begin{aligned}
P_1 &= \frac{K_1 C D_0 J}{(1 + K_1 J + K_3 M)} \\
P_2 &= \frac{K_3 C D_0 M}{(1 + K_1 J + K_3 M)} \\
P_3 &= \frac{K_5 X Y_0 M}{(1 + K_5 M + K_7 J)} \\
P_4 &= \frac{K_7 X Y_0 J}{(1 + K_5 M + K_7 J)}.
\end{aligned}
\tag{15}
$$

*J* and *M* also fulfill that they are either free or forming efflux pumps, allowing to define total concentrations:

$$
\begin{aligned}
J_0 &\equiv J + P_1 + P_4 \\
M_0 &\equiv M + P_2 + P_3.
\end{aligned}
\tag{16}
$$

Substituting *J* and *M* by $J_0$ and $M_0$ in equations (15):

$$
\begin{aligned}
P_1 &= \frac{K_1 C D_0 (J_0 - P_1 - P_4)}{1 + K_1 (J_0 - P_1 - P_4) + K_3 (M_0 - P_2 - P_3)} \\
P_2 &= \frac{K_3 C D_0 (M_0 - P_2 - P_3)}{1 + K_1 (J_0 - P_1 - P_4) + K_3 (M_0 - P_2 - P_3)} \\
P_3 &= \frac{K_5 X Y_0 (M_0 - P_2 - P_3)}{1 + K_5 (M_0 - P_2 - P_3) + K_7 (J_0 - P_1 - P_4)} \\
P_4 &= \frac{K_7 X Y_0 (J_0 - P_1 - P_4)}{1 + K_5 (M_0 - P_2 - P_3) + K_7 (J_0 - P_1 - P_4)}.
\end{aligned}
\tag{17}
$$

In the case where the limiting factor to build efflux pumps is the concentration of OprJ and OprM, we can assume that $J_0$ and $M_0$ are basically all sequestered into efflux pumps, therefore:

$$
\begin{aligned}
(J_0 - P_1 - P_4) &\ll 1 \\
(M_0 - P_2 - P_3) &\ll 1,
\end{aligned}
\tag{18}
$$

and we can write the approximate equations:

$$
\begin{aligned}
P_1 &\approx K_1 CD_0 (J_0 - P_1 - P_4) \\
P_2 &\approx K_3 CD_0 (M_0 - P_2 - P_3) \\
P_3 &\approx K_5 XY_0 (M_0 - P_2 - P_3) \\
P_4 &\approx K_7 XY_0 (J_0 - P_1 - P_4).
\end{aligned} \tag{19}
$$

This is a linear system of 4 equations and 4 variables that we can solve:

$$
\begin{aligned}
P_1 &= \frac{K_1 CD_0}{1 + K_1 CD_0 + K_7 XY_0} J_0 \\
P_2 &= \frac{K_3 CD_0}{1 + K_3 CD_0 + K_5 XY_0} M_0 \\
P_3 &= \frac{K_5 XY_0}{1 + K_3 CD_0 + K_5 XY_0} M_0 \\
P_4 &= \frac{K_7 XY_0}{1 + K_1 CD_0 + K_7 XY_0} J_0
\end{aligned} \tag{20}
$$

If any $K_i$ is zero, the solution is still well-defined: $K_7 = 0$ would imply $P_4 = 0$, as assummed in the Main Text. $P_1$ depends linearly on $CD_0$ when $CD_0 \ll K_1^{-1}$, and then eventually saturates as $CD_0$ increases. $P_1$ decreases monotonically with $XY_0$. Similar considerations apply to the other efflux pumps. The concentration of MexCD-containing efflux pumps is:

$$
P_1 + P_2 = \frac{K_1 CD_0}{1 + K_1 CD_0 + K_7 XY_0} J_0 + \frac{K_3 CD_0}{1 + K_3 CD_0 + K_5 XY_0} M_0 \tag{21}
$$

This is a monotonically increasing function of [*MexCD*]. From equations (20) and (21) we derive the conclusions presented in the Main Text: the concentration of MexCD-containing efflux pumps ($P_1$+$P_2$) increases monotonically with the total MexCD concentration, and the concentration of MexXY-containing efflux pumps (be it $P_3$ if $K_7 = 0$, equation (2) of the Main Text, or $P_3 + P_4$ otherwise) decreases with MexCD concentration. Since *mexCD* expression depends quadratically on DC, equation (10), we can write efflux pump concentrations as a function of [$DC_{out}$] just substituting in equations (20) $CD_0$ with the [$DC_{out}$] formula given in equation (10).

We can see that the solution equations (20) is consistent with the approximation in equations (18). This approximation should be valid for $K_i \gg 1$, and in that case equations (20) can be approximated by:

$$\begin{aligned} P_1 &= \frac{K_1 CD_0}{K_1 CD_0 + K_7 XY_0} J_0 \\ P_2 &= \frac{K_3 CD_0}{K_3 CD_0 + K_5 XY_0} M_0 \\ P_3 &= \frac{K_5 XY_0}{K_3 CD_0 + K_5 XY_0} M_0 \\ P_4 &= \frac{K_7 XY_0}{K_1 CD_0 + K_7 XY_0} J_0 \end{aligned} \tag{22}$$

These expressions result in $P_1 + P_4 = J_0$ and $P_2 + P_3 = M_0$, consistent with the approximation made.

The assumption was made that OprJ and OprM were the limiting species. The prediction for their mutants under this scenario would be that their respective efflux pumps do not form, but the other efflux pumps are not affected. Mutations in MexCD or MexXY, however, would have a positive effect on the efflux pumps where they are not components.

**Supplementary Note 3: relationship between amount of efflux pumps and MIC**

The concentration of antibiotic $A$ inside the cell follows this equation, modified from [2]:

$$\dot{A} = \phi_{\text{in}} A_{\text{out}} - \phi_{\text{out}} A - \frac{v P^n}{K_M + P^n} A - \lambda A,$$

where $A_{out}$ is the concentration of antibiotic in the medium outside the cell, $\phi_{in}$ and $\phi_{out}$ are the diffusion constants for the antibiotic in and out of the cell, respectively, $v$ is the maximal efflux velocity, $K_M$ is the Michaelis constant for the efflux pump $P$ (either MexCD-OprJ or MexXY-OprM), $n$ is the cooperativity of the efflux pump and $\lambda$ is bacterial growth rate. The intracellular antibiotic concentration can increase from inflow of external antibiotic, and can decrease from outflow, efflux or dilution due to cell division.

Now, assuming steady state ($\dot{A} = 0$), we have:

$$\phi_{\mathrm{in}} A_{\mathrm{out}} = \left( \phi_{\mathrm{out}} + \frac{vP^n}{K_M + P^n} + \lambda \right) A. \tag{23}$$

Our experimental data gives us the minimal inhibitory concentration (MIC) for two antibiotics, ciprofloxacin (CIP) and tobramycin (TOB), across several DC concentrations. At the MIC, growth ceases, $\lambda \to 0$. To understand the corresponding intracellular antibiotic concentration *A*, we must consider how each antibiotic inhibits growth.

*Ciprofloxacin:* CIP inhibits DNA gyrase, an essential enzyme for DNA replication and transcription. Following a simplified mechanistic model (which could be refined using the framework of [3], we assume growth rate is proportional to functional gyrase concentration, which decreases with intracellular CIP. This yields:

$$\lambda = \frac{\lambda_0}{1 + k_{in} A^k}$$

where $\lambda_0$ is the maximum growth rate in the absence of antibiotic, $k$ = 1, 2, 3,... reflects the cooperativity of CIP binding to gyrase, and $k_{in}$ is an effective inhibition constant.

*Tobramycin*: TOB binds the ribosome, inhibiting protein synthesis. Greulich et al. [2] developed a detailed model linking ribosome-binding antibiotics to bacterial growth using the growth laws of Scott and Hwa [4]. From their framework, we obtain:

$$\lambda = \frac{\lambda_0}{1 + k_{in}A}$$

where $k_{in}$ can be expressed in terms of the binding kinetics and ribosome parameters:

$$k_{in} = 4\frac{k_{\text{on}}\lambda_0}{k_{\text{off}}\kappa_t\Delta R}.$$

Here, $k_{\text{on}}$ and $k_{\text{off}}$ are the association and dissociation rate constants for TOB-ribosome binding, $\kappa_t$ is the translation elongation rate, and ΔR represents the ribosome pool available for antibiotic binding.

In both cases, growth rate $\lambda$ depends inversely on intracellular antibiotic concentration $A$. When $\lambda \to 0$ as $A_{out} \to MIC$, mathematically, $A \to \infty$. However, in practice, growth becomes undetectable when $\lambda$ falls below a given threshold. Setting $\epsilon = \lambda/\lambda_0 \ll 1$ as this threshold, we obtain:

$$1 + k_{in}A = \frac{1}{\epsilon} \Longrightarrow A = \frac{\frac{1}{\epsilon} - 1}{k_{in}} \approx \frac{1}{\epsilon k_{in}}. \tag{24}$$

Note that if $k > 1$ in the CIP case, equation (24) becomes $A \approx (\epsilon k_{in})^{-1/k}$, but since both $\epsilon$ and $k_{in}$ are unknown, we can absorb this into the parameterization without loss of generality.

Going back to equation (23), taking the limit $A_{out} \to MIC$ and substituting the approximation in (24), we obtain

$$\phi_{\text{in}}\text{MIC} \approx \left(\phi_{\text{out}} + \frac{vP^n}{K_M + P^n}\right)\frac{1}{\epsilon k_{in}}\text{MIC} \approx \frac{\phi_{\text{out}}}{\epsilon k_{in}\phi_{in}} + \frac{v}{\epsilon k_{in}\phi_{in}}\frac{P^n}{K_M + P^n},$$

but since most of these are unknown parameters for us, we can rewrite this expression as:

$$\text{MIC} \approx a + b\frac{P^n}{K_M + P^n}. \quad (25)$$

Now, we are interested in studying how the MIC to CIP and TOB will change as we increase the concentration of DC. Equation (25) shows that the MIC is a monotonic increasing function of the concentration of the efflux pumps involved. For CIP, we consider that those efflux pumps are the MexCD–OprJ and MexCD–OprM complexes, and equation (21) shows that in this case the concentration *P* is an increasing function of [*MexCD*], and therefore so is the MIC. Finally, in Figure 1 we see that [*MexCD*] increases with [*DC*]. Therefore, equation (25) predicts that the MIC to CIP is a monotonic increasing function of [*DC*]. Conversely, the *P* in equation (25) for the MIC to TOB is the concentration of the MexXY–OprM complex, that is a monotonically decreasing function of [*MexCD*], as seen in the formula for $P_3$ in equations (20), and therefore the MIC to TOB is a decreasing function of [*DC*]. Assuming full promiscuity and that a MexXY–OprJ complex could also form and pump out TOB would not change this conclusion, since MexXY–OprJ is also a decreasing function of [*MexCD*] as seen in the formula for $P_4$ in equation (20), and therefore $P_3$+$P_4$ is also a decreasing function of [*MexCD*] or [*DC*].

To make these conclusions quantitative, we have to take the formula for [*MexCD*] dependence on [*DC*], equation (10), substitute it in the equations (20) for the relevant efflux pump concentrations, and substitute those in equation (25) for the corresponding MIC. Assuming for simplicity $n$=1 in equation (25) (the fits in Figure 3 show that this is consistent with the data), for CIP this results in

$$\mathrm{MIC_{CIP}} = a_{CIP} + b_{CIP} \frac{P_1([DC]) + P_2([DC])}{K_{M,CIP} + P_1([DC]) + P_2([DC])}. \tag{26}$$

We have written $\mathrm{MIC_{CIP}}$ as a ratio of quadratic polynomials in $[DC]^2$. This form introduces additional higher-order coefficients (the $[DC]^4$ terms) that are typically poorly identifiable from finite, noisy MIC–DC datasets because they are strongly correlated with lower-order terms (baseline, dynamic range, and crossover scale) [5]. For this reason, we use a parsimonious surrogate that preserves the correct qualitative behavior (monotonic increase and saturation with $[DC]^2$) while reducing the number of free parameters:

$$MIC_{CIP} = A_{CIP} \frac{1 + C_{CIP}\left(\frac{[DC]}{B_{CIP}}\right)^2}{1 + \left(\frac{[DC]}{B_{CIP}}\right)^2}. \tag{27}$$

Equation (27) captures the three key features that are robustly constrained by the data: baseline MIC at low DC, saturation MIC at high DC, and the crossover scale in $[DC]$. The neglected higher-order terms correspond to additional curvature that is difficult to estimate reliably without substantially denser datasets or strong priors.

For TOB, similar considerations apply, but if we actually only consider the MexXY–OprM efflux pump, then:

$$MIC_{TOB} = a_{TOB} + b_{TOB} \frac{P_3([DC])}{K_{M,TOB} + P_3([DC])}. \tag{28}$$

Defining,

$$N \equiv M_0 K_5 X Y_0, \qquad D_0 \equiv 1 + K_5 X Y_0, \qquad p \equiv N + K_{M,TOB} D_0, \qquad q \equiv a K_{M,TOB} K_3 \tag{29}$$

and

$$A_{TOB} \equiv a_{TOB} + b_{TOB} \frac{N\,c}{pc+q}, \qquad B_{TOB} \equiv b\sqrt{\frac{pc+q}{p+q}}, \qquad C_{TOB} \equiv 1 + \frac{b_{TOB} N q (1-c)}{(p+q)[a_{TOB}(pc+q) + b_{TOB} N c]}, \tag{30}$$

we can write $MIC_{TOB}$ as

$$MIC_{TOB} = A_{CIP} \frac{1 + C_{TOB}\left(\frac{[DC]}{B_{TOB}}\right)^2}{1 + \left(\frac{[DC]}{B_{TOB}}\right)^2}. \tag{31}$$

Now we do not need to make any ad hoc simplifications as we did with CIP, since we already have only three free parameters to capture the shape of MIC decay with DC. Note that even if we considered a fully promiscuous model where a MexXY–OprJ complex could also form and pump out TOB, equation (31) would be a parsimonious surrogate that preserves the correct qualitative behavior (monotonic decrease over a finite DC scale of [*DC*]) to reduce the number of free parameters, therefore being a convenient fit function. Also note that the functional forms for $MIC_{CIP}$ and $MIC_{TOB}$ are the same, but since the parameter $c$>1 (Figure 1B), meaning that *[MexCD]* increases with *[DC]*, then $C_{TOB}$ defined in equation (30) is smaller than 1, and therefore $MIC_{TOB}$ is a decreasing function of *[DC]*.

Equations (27) and (31) are equation (3) in the Main Text, that we fit to experimental data (see Bayesian estimation). The mean and standard deviation obtained from the posterior distribution of the fit parameters are: $A_{CIP}$ =0.07±0.02, $B_{CIP}$ =0.39±0.37, $C_{CIP}$ =2.1±0.6, $A_{TOB}$ =0.22±0.04, $B_{TOB}$ =0.3±0.3, $C_{TOB}$ =0.4±0.2. While the values determining baseline MIC at low DC and saturation MIC at high DC are relatively well determined by parameters *A* and *C*, the data constrains poorly the crossover scale, since the *B* parameters are not well determined.

| name | average (kcal/mol) | vina (kcal/mol) | nnscore (kcal/mol) | dlscore (kcal/mol) | average (pKd) | vina (pKd) | nnscore (pKd) | dlscore (pKd) |
|---|---|---|---|---|---|---|---|---|
| **RamR/vina/RamR_berberine_1** | **-9.26** | **-9.49** | **-9.10** | **-9.19** | **5.30** | **2.43** | **6.70** | **6.77** |
| **RamR/vina/RamR_chenodeoxycolic_1** | **-9.65** | **-9.48** | **-9.05** | **-10.43** | **5.59** | **2.43** | **6.66** | **7.68** |
| **RamR x dequalinium** | **-8.31** | **-7.69** | **-9.76** | **-7.47** | **4.88** | **1.97** | **7.19** | **5.50** |
| **RamR x rhodamine6G** | **-7.39** | **-8.51** | **-6.13** | **-7.54** | **4.08** | **2.18** | **4.51** | **5.55** |
| NfxB x DC 06 | -10.36 | -8.48 | -12.38 | -10.21 | 6.27 | 2.17 | 9.11 | 7.51 |
| NfxB x DC 08 | -9.79 | -8.58 | -9.52 | -11.26 | 5.83 | 2.20 | 7.01 | 8.29 |
| NfxB x DC 01 | -9.75 | -8.65 | -9.89 | -10.72 | 5.80 | 2.22 | 7.28 | 7.89 |
| NfxB x DC 02 | -9.68 | -8.79 | -9.62 | -10.64 | 5.72 | 2.25 | 7.08 | 7.83 |
| NfxB x DC 07 | -9.51 | -8.24 | -9.81 | -10.47 | 5.68 | 2.11 | 7.22 | 7.71 |
| NfxB x DC 05 | -9.40 | -8.76 | -8.73 | -10.72 | 5.52 | 2.24 | 6.43 | 7.89 |
| NfxB x DC 03 | -9.25 | -8.75 | -8.25 | -10.74 | 5.41 | 2.24 | 6.07 | 7.91 |
| NfxB x DC 04 | -9.05 | -8.22 | -8.10 | -10.82 | 5.35 | 2.11 | 5.97 | 7.97 |
| NfxB x DC 09 | -8.94 | -7.86 | -8.35 | -10.61 | 5.32 | 2.01 | 6.15 | 7.81 |
| NfxB x DC 10 | -8.19 | -7.53 | -6.49 | -10.56 | 4.83 | 1.93 | 4.78 | 7.77 |

**Supplementary Table 1.** Affinity values of RamR for dequalinium chloride (DC) and rhodamine 6G, and of NfxB for DC, calculated using various scoring programs: RamR affinities were obtained using cognate conformations to be used as a reference. NfxB affinities were calculated for the ten best conformations obtained after docking the SwissModel-derived model of NfxB with DC with Autodock Vina and are shown as "NfxB x DC" followed by the conformation number and sorted by average affinity score. Since no score has proven systematically more reliable, we have used Autodok Vina, NNscore, and DLscore. Values are shown as both kcal/mol (more negative means more energy is released on binding, i.e. it is more favorable) and pKd (negative base-10 logarithm of the dissociation constant, $pK_d$, higher values imply higher affinity). Scores were averaged to calculate a consensus vote estimation across methods of the affinity.

| DC | CIPm | CIPM | TOBm | TOBM | CIPavg | TOBavg |
|---|---|---|---|---|---|---|
| 0 | 0.035 | 0.05 | 0.2 | 0.25 | 0.0425 | 0.225 |
| 0.1 | 0.07 | 0.1 | 0.2 | 0.25 | 0.085 | 0.225 |
| 0.2 | 0.07 | 0.1 | 0.1 | 0.12 | 0.085 | 0.11 |
| 0.3 | 0.1 | 0.12 | 0.1 | 0.12 | 0.11 | 0.11 |
| 0.4 | 0.12 | 0.16 | 0.1 | 0.12 | 0.14 | 0.11 |
| 0.5 | 0.12 | 0.16 | 0.1 | 0.12 | 0.14 | 0.11 |
| 0.8 | 0.12 | 0.16 | 0.1 | 0.12 | 0.14 | 0.11 |
| 1 | 0.12 | 0.16 | 0.1 | 0.12 | 0.14 | 0.11 |

**Supplementary Table 2**. Minimal inhibitory concentrations (MICs, µg/mL) of ciprofloxacin (CIP) and tobramycin (TOB) across dequalinium chloride (DC) concentrations. CIPm/TOBm indicate the highest concentrations permitting growth, CIPM/TOBM the lowest preventing growth, and CIPavg/TOBavg their midpoints.

### REFERENCES


1 Laborda, P., Alcalde-Rico, M., Blanco, P., Martínez, J. L. & Hernando-Amado, S. Novel Inducers of the Expression of Multidrug Efflux Pumps That Trigger Pseudomonas aeruginosa Transient Antibiotic Resistance. *Antimicrobial agents and chemotherapy* **63**, doi:10.1128/aac.01095-19 (2019).

2 Greulich, P., Scott, M., Evans, M. R. & Allen, R. J. Growth-dependent bacterial susceptibility to ribosome-targeting antibiotics. *Molecular systems biology* **11**, 796, doi:10.15252/msb.20145949 (2015).

3 Ojkic, N. *et al.* A Roadblock-and-Kill Mechanism of Action Model for the DNA-Targeting Antibiotic Ciprofloxacin. *Antimicrobial agents and chemotherapy* **64**, doi:10.1128/AAC.02487-19 (2020).

4 Scott, M., Gunderson, C. W., Mateescu, E. M., Zhang, Z. & Hwa, T. Interdependence of cell growth and gene expression: origins and consequences. *Science* **330**, 1099-1102, doi:10.1126/science.1192588 (2010).

5 Massonis, G., Villaverde, A. F. & Banga, J. R. Distilling identifiable and interpretable dynamic models from biological data. *PLoS computational biology* **19**, e1011014, doi:10.1371/journal.pcbi.1011014 (2023).
